# Aβ*56 is present in brain extracts enriched for extracellular proteins

**DOI:** 10.1101/2024.10.09.617424

**Authors:** Peng Liu, Ian P. Lapcinski, Karen H. Ashe

## Abstract

Amyloid-β (Aβ) oligomers are believed to be important in the pathogenesis of Alzheimer’s disease (AD). Aβ*56 is an Aβ oligomer that has been reported in several lines of transgenic mice modeling AD, including Tg2576, hAPP-J20, 3xTgAD, 5xFAD, and APPTTA. In Tg2576 mice, Aβ*56 appears several months before neuritic plaques and impairs memory when injected into the brains of healthy rodents. Aβ*56 has several distinctive biochemical features: 1) it is soluble in aqueous buffers, 2) it is stable in the ionic detergent sodium dodecyl sulfate (SDS), 3) it has an apparent mass of 56 kDa whether measured by denaturing SDS polyacrylamide gel electrophoresis or non-denaturing size-exclusion chromatography, 4) it binds to A11 conformational antibodies that recognize non-fibrillar oligomeric assemblies, and 5) it contains canonical Aβ(1-40) and/or Aβ(1-42). Here, we show its presence in Tg2576 brain extracts enriched for extracellular proteins and conclude that Aβ*56 is present in the extracellular space.

## Introduction

Amyloid-β (Aβ) oligomers are believed to be important in the pathogenesis of Alzheimer’s disease (AD). Aβ*56 is an Aβ oligomer that has been described in papers from several labs^1–11^ and has been found in at least five lines of transgenic AD mouse models, including Tg2576^7,11^, hAPP-J20^5^, 3xTgAD^1^, 5xFAD^11^, and APPTTA^11^. In Tg2576 mice, Aβ*56 appears several months before neuritic plaques, which form at 12-13 months, but has not been detected before 6 months of age^11^. Aβ*56 isolated from Tg2576 mice impairs memory when injected into the brains of healthy rodents^11^. Aβ*56 has been shown to correspond to memory loss better than neuritic plaques in Tg2576^7,8,11^, hAPP-J20^5,6^, and APPTTA mice^9^.

Aβ*56 has several distinctive biochemical features: 1) it is soluble in aqueous buffers^11^, 2) it is stable in the ionic detergent sodium dodecyl sulfate (SDS)^11^, 3) it has an apparent mass of 56 kDa whether measured by denaturing SDS-polyacrylamide gel electrophoresis (SDS-PAGE) or non-denaturing size-exclusion chromatography (SEC)^11^, 4) it binds to A11 conformational antibodies that recognize non-fibrillar oligomeric assemblies^11^, and 5) it contains canonical Aβ(1-40) and/or Aβ(1-42)^11^. No other known high-molecular-weight, brain-derived Aβ oligomer exhibits these biochemical features. In particular, the stability of a high-molecular-weight oligomer in SDS is unique to Aβ*56.

In a recent paper, we confirmed the existence of Aβ*56 in aqueous brain extracts but did not specifically demonstrate its presence in the extracellular space, since we extracted Aβ*56 using an aqueous buffer that does not differentiate between extracellular and non-extracellular proteins^11^. In the current study, we validated the effectiveness a procedure that compartmentalizes extracellular and membrane-associated proteins and, using that procedure, found that Aβ*56 is present in extracts enriched for extracellular proteins.

## Results

We validated a method to compartmentalize a set of known cellular proteins into their respective extracellular and membrane-associated compartments. We used western blotting (WB) depicted in **Fig. 1a** to determine the fidelity of this fractionation process. The results show that proteins in the extracellular extracts are enriched for the extracellular protein soluble amyloid precursor protein-α (sAPPα) (**Fig. 1b**) and those in the membrane extracts are enriched for the membrane-associated protein, full-length amyloid precursor protein (fl-APP) (**Fig. 1c**), confirming that the compartmentalization procedure effectively enriches extracts for extracellular and membrane-associated proteins accordingly. We named extracts that are enriched for extracellular proteins “EC” and extracts that are enriched for membrane-associated proteins “MB.”

**Figure 1.**
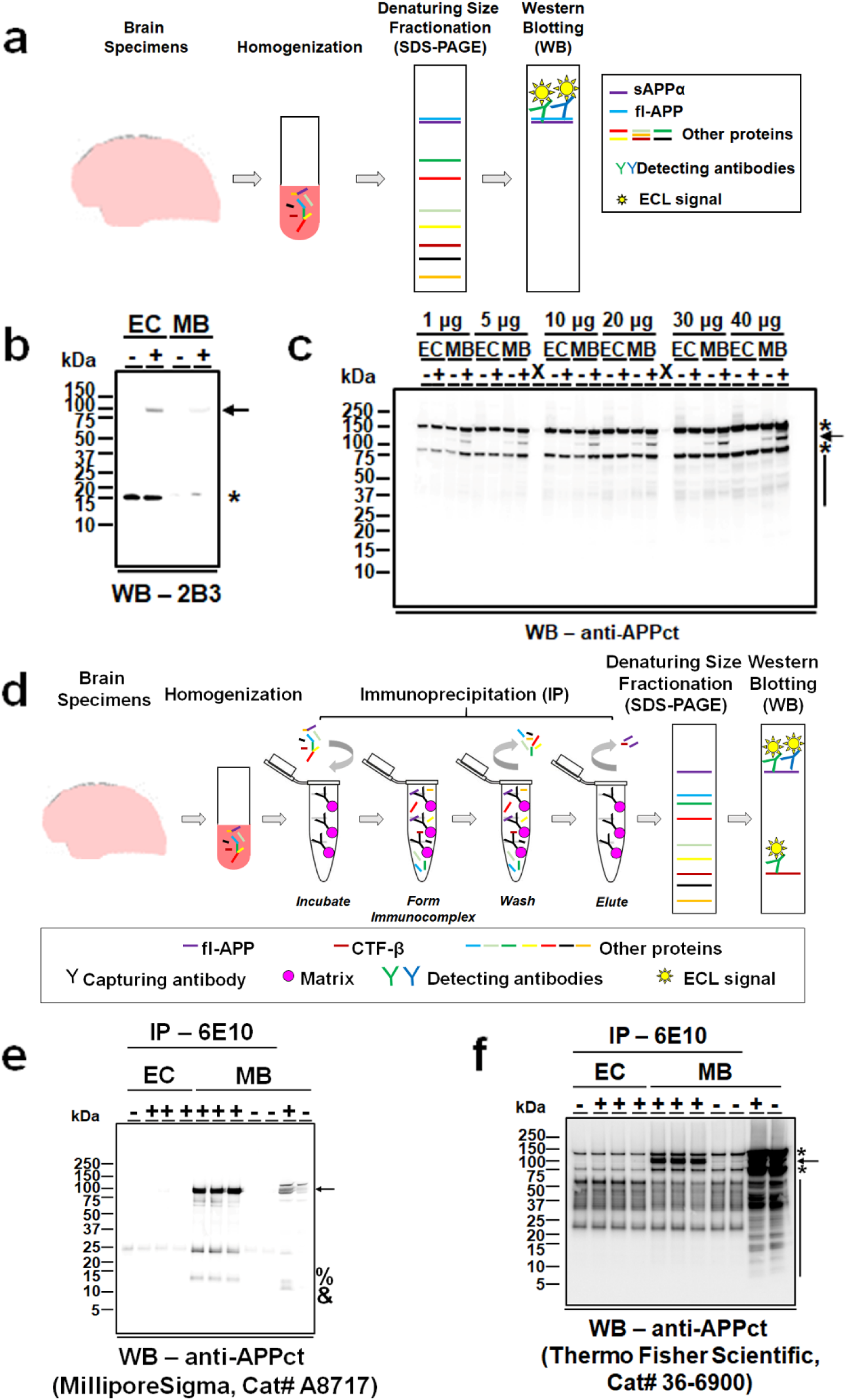
No full-length APP is detected in EC of Tg2576 mice modeling AD. **a**) A schematic diagram illustrating the experimental procedure of figures b and c. Briefly, forebrain tissue was subjected to homogenization in detergent-containing buffers to obtain EC and MB. Proteins of varied sizes were then subjected to SDS-PAGE under denaturing conditions, and WB was then carried out to detect sAPPα (b) and full-length APP (fl-APP) (c) using detecting antibodies coupled to ECL-based signal detection. Detailed descriptions were provided in the *Method* section. **b**) A representative WB image showing the preferential detection of sAPPα (arrow) in the EC of Tg2576 mice relative to the MB using the monoclonal 2B3 antibody (IBL America, Cat# 11088) that recognizes specifically Aβ or APP forms ending C-terminally at amino acid 15 of Aβ. +, combined EC or MB of thee 9-month, female Tg2576 mice at a volume ratio of 1:1:1; -, combined EC or MB of two age-matched, female, non-transgenic littermates of Tg2576 mice at a volume ratio of 1:1. For each lane, brain extracts containing 85 µg of total proteins were used. **c**) A representative WB image showing the preferential detection of fl-APP (arrow) in the MB of Tg2576 mice relative to the EC using a polyclonal antibody (anti-APPct; Thermo Fisher Scientific, Cat# 36-6900) that recognizes specifically the C-terminus of fl-APP. The fl-APP detection pattern in the EC and MB was examined at six different levels of total proteins (1, 5, 10, 20, 30, and 40 µg). +, combined brain extracts of thee 9-month, female Tg2576 mice at a volume ratio of 1:1:1; -, combined brain extracts of two age-matched, female, non-transgenic littermates of Tg2576 mice at a volume ratio of 1:1. X, empty lane. **d**) A schematic diagram illustrating the experimental procedure of figures e and f. Briefly, forebrain tissue was subjected to homogenization in detergent-containing buffers to obtain EC and membrane-bound brain extracts (MB). Proteins of varied sizes were then subjected to immunoprecipitation (IP) using the 6E10 capturing antibody bound to matrix. Next, the 6E10-precipitated entities were eluted and size-fractionated by SDS-PAGE under denaturing conditions, and WB was then carried out using detecting antibodies coupled to ECL-based signal detection. Detailed descriptions were provided in the *Method* section. **e**) Representative IP/WB analyses showing the detection of the fl-APP (arrow) and C-terminal APP fragment processed by β-secretase (CTF-β, %) in the 6E10-precipitated MB but not EC of Tg2576 mice using a polyclonal antibody (anti-APPct; MilliporeSigma, Cat# A8717) that recognizes specifically the C-terminus of fl-APP. Besides fl-APP and CTF-β, C-terminal APP fragment processed by α-secretase (CTF-α, &) is also detected in the MB by straight WB. **f**) Representative IP/WB analyses showing the detection of the fl-APP in the 6E10-precipitated MB but not EC of Tg2576 mice using a polyclonal antibody (anti-APPct; Thermo Fisher Scientific, Cat# 36-6900) that recognizes specifically the C-terminus of fl-APP. For e and f, +, brain extracts of 9-month, female Tg2576 mice; -, brain extracts of age-matched, female, non-transgenic littermates of Tg2576 mice. Brain extracts containing 100 µg of total proteins were used for IP/WB in each lane, and brain extracts containing 40 µg of total proteins in the MB were used for straight WB in each lane. For b, c, e, and f, * and vertical line, non-specific bands; kDa, kilo-Daltons.

We also showed that this technique compartmentalizes the membrane-associated C-terminal fragment (CTF)-β APP appropriately. As depicted in **Fig. 1d**, we immunoprecipitated proteins from EC or MB using 6E10 antibodies, which recognize amino-acids 3-8 in Aβ, and identified proteins containing the C-terminus of APP by WB using anti-APPct antibodies from two different vendors (**Figs. 1e** and **1f**). MilliporeSigma 6E10 antibodies readily detected fl-APP along with CTF-β in MB but not EC (**Fig. 1e**), while Thermo Fisher Scientific 6E10 antibodies detected fl-APP in MB, but obscured CTF-β due to non-specific binding and/or lower binding affinity (**Fig. 1f**).

Next, we established the presence of Aβ*56 in EC extracts. As depicted in **Fig. 2a**, EC from 9-month-old, neuritic-plaque-free Tg2576 mice were fractionated by SEC under detergent-free conditions and detected by WB after fractionation by SDS-PAGE using 82E1 antibodies recognizing the N-terminus neo-epitope of Aβ(1-x). The upper panel of **Fig. 2b** shows a ∼56-kDa band peaking in fractions 22-23 corresponding to ∼40-56 kDa (**Fig. 2c**). The middle panel in **Fig. 2b** shows a higher exposure of the portion of the blot containing monomers. The lower panel of **Fig. 2b** shows non-specific signals identified prior to antibody exposure. These results indicate that the 82E1-reactive entity in EC extracts migrates at ∼56 kDa under both non-denaturing and denaturing conditions.

**Figure 2.**
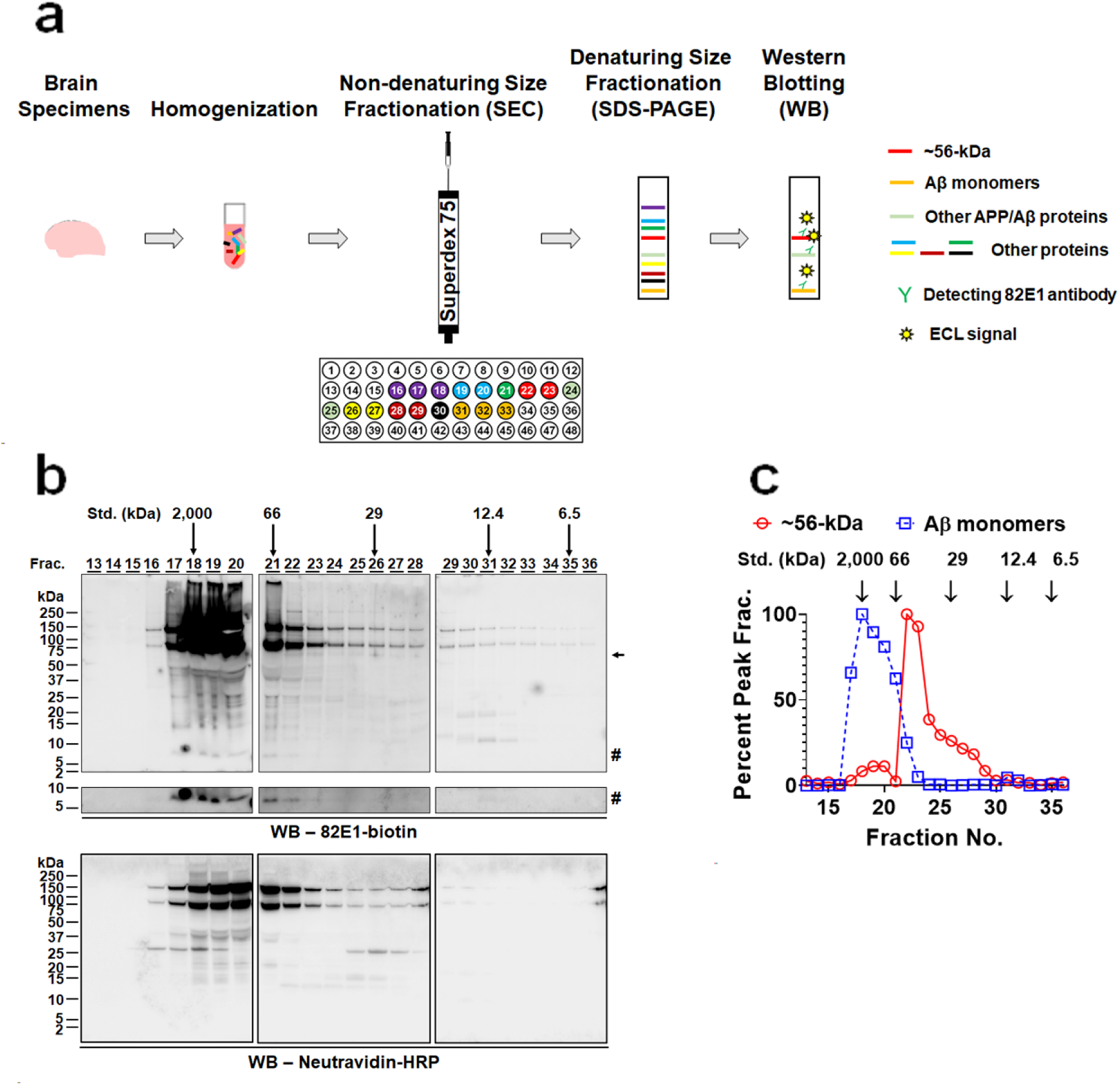
A ∼56-kDa, 82E1-reactive entity present in the brain of Tg2576 mice modeling AD is not artificially formed by exposure to SDS. **a**) A schematic diagram illustrating the experimental procedure. Briefly, forebrain tissue was subjected to homogenization in a detergent-containing buffer to obtain EC. Proteins of varied sizes were then separated under non-denaturing conditions by SEC using a Superdex 75 column and collected into a total of 48 fractions. Next, proteins in fractions of interest were subjected to SDS-PAGE under denaturing conditions, and WB was then carried out to detect APP) metabolites and Aβ entities using monoclonal antibody 82E1 that recognizes the N-terminus of Aβ (1-x) coupled to electrochemiluminescence (ECL)-based signal detection. Detailed descriptions were provided in the *Method* section. **b**) Representative WB analyses showing the detection of a ∼56-kDa entity (arrow) and Aβ monomers (#) in SEC fractions 13-36 of EC of 9-month Tg2576 mice using the biotinylated 82E1 (82E1-biotin) antibody. The three WB images (*i.e.*, Frac. 13-20, 21-28, and 29-36) shown were obtained using the same exposure time (*Upper panels*, 10 sec). To better visualize Aβ monomers, WB images obtained using a longer exposure time are also shown (*Middle panels*, 100 sec exposure). To differentiate the 82E1-specific signals from those unrelated to the primary antibody, WB was initially probed with horseradish peroxidase (HRP)-conjugated Neutravidin (Neutravidin-HRP; *lower panels*, 100 sec exposure). Neither the ∼56-kDa entity nor Aβ monomers are detected. **c**) Quantitative analyses of levels of the ∼56-kDa entity and Aβ monomers in SEC fractions. The level of the ∼56-kDa entity in each fraction is normalized to its highest level in fraction 22; and Aβ monomers, fraction 18. The elution profile of biomolecule standards (Std.) of varied sizes is shown above the WB images. kDa, kilo-Daltons.

Quantitative analyses showed two distinct peaks separating the ∼56-kDa entity from monomers (**Fig. 2c**), confirming that it was not generated artificially due to exposure to SDS, which has been shown to promote Aβ aggregation under some experimental conditions^12–14^.

Finally, we showed that the ∼56 kDa entity in EC is a non-fibrillar oligomer using the methodology depicted in **Fig. 3a**. We immunoprecipitated proteins eluting in SEC fractions corresponding to ∼56 kDa using D8Q7I recognizing the C-terminus neo-epitope of Aβ(x-40). Next, we fractionated the immunoaffinity-purified proteins by SDS-PAGE, excised and electro-eluted the ∼56-kDa entity, and probed a WB of the electro-eluted material with A11 polyclonal antibodies recognizing epitopes present in non-fibrillar, but not fibrillar, oligomers ^15^. **Fig. 3b** shows that the ∼56 kDa D8Q7I-reactive entity binds A11 antibodies. In parallel, EC from age-matched non-transgenic littermates of Tg2576 mice were similarly processed, and the A11 antibodies showed no reactivity to the electro-eluted proteins (**Fig. 3b**).

**Figure 3.**
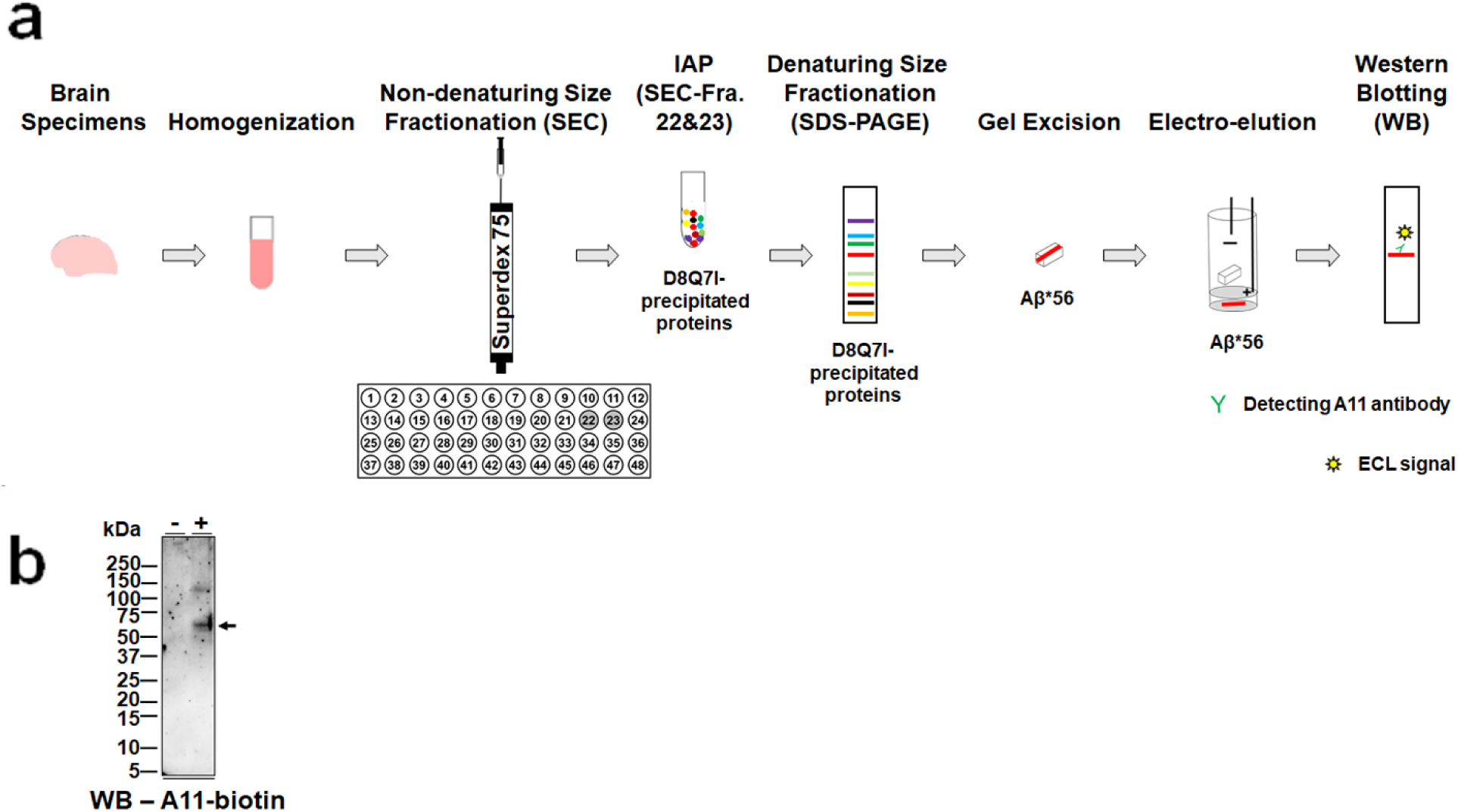
The ∼56-kDa, 82E1-reactive entity present in the EC of Tg2576 mice modeling AD is captured by an anti-Aβ antibody and reactive to the anti-oligomer A11 antibody. **a**) A schematic diagram illustrating the experimental procedure. Briefly, forebrain tissue was subjected to homogenization in a detergent-containing buffer to obtain EC. Proteins of varied sizes were then separated under non-denaturing conditions by SEC using a Superdex 75 column and collected into a total of 48 fractions. Next, proteins in fractions 22-23 where the ∼56-kDa, 82E1-reactive entity was enriched were subjected to immunoaffinity purification (IAP) by the rabbit monoclonal antibody D8Q7I that is directed against the C-terminus of Aβ(x-40). The captured proteins were eluted and then size-fractionated by SDS-PAGE. The ∼56-kDa entity was then separated from other D8Q7I-precipiated proteins and released from polyacrylamide gel via electro-elution. Finally, the purified ∼56-kDa entity was probed in semi-denaturing WB analysis by rabbit polyclonal A11 antibody to validate it as an oligomeric assembly. Of note, EC prepared from age- and gender-matched, non-transgenic littermates of Tg2576 mice were subjected to the same experimental procedures, which serve as a negative control. Detailed descriptions were provided in the *Method* section. **b**) Representative WB analyses (60 sec exposure) showing the detection of a ∼56-kDa entity (arrow) purified from EC of Tg2576 mice (+) but not their non-transgenic littermates (-) using the biotinylated anti-oligomer A11 (A11-biotin) antibody. kDa, kilo-Daltons.

Taken altogether, the results in **Figs. 1-3** show that brain extracts enriched for extracellular proteins contain an entity with the key biochemical properties of Aβ*56, namely that it is SDS-stable, A11-reactive, and has a molecular mass of ∼56 kDa. We concluded that the extracellular space contains Aβ*56.

## Discussion

In this paper, we demonstrated that it is possible to biochemically separate extracellular from membrane-associated proteins. Using this methodology, we identified an entity in fractions enriched for extracellular proteins that is stable in SDS, contains canonical Aβ, binds A11 antibodies that uniquely recognize non-fibrillar oligomers, and corresponds to ∼56 kDa by both non-denaturing SEC and denaturing SDS-PAGE. Since Aβ*56 is the only known brain-derived Aβ assembly to possess these biochemical properties, the results indicate that the extracellular space contains Aβ*56.

The results in **Fig. 1** confirm the ability to compartmentalize fl-APP and its cleavage products, CTF-β and sAPPα, into known cellular compartments. Fl-APP and CTF-β appropriately appear in membrane-enriched but not extracellular-enriched fractions (**Figs. 1b, c, f**), and sAPPα appropriately appears in extracellular-enriched but not membrane-enriched fractions (**Fig. 1e**).

The results in **Figs. 2** and **3** confirm the existence of a ∼56-kDa Aβ oligomer in the extracellular space. The Aβ molecules in this oligomer did not aggregate artificially due to exposure to SDS (**Fig. 2**) and are arranged in the manner of non-fibrillar oligomers (**Fig. 3**), though precisely how the molecules are arranged remains unknown.

Aβ*56 was initially identified 18 years ago in a paper by Lesné *et al.*^16^. This study described Aβ*56 as a soluble, SDS-stable, extracellular Aβ oligomer that binds A11 antibodies, impairs memory, and has a molecular mass of ∼56 kDa whether measured by denaturing SDS-PAGE or non-denaturing SEC. Lesné *et al.*^16^ proposed that oligomers were the primary form of Aβ in the brain responsible for memory impairment, addressing the previously unclear question of whether monomers, oligomers or fibrils were the main contributors. Since then, several other types of Aβ oligomers have been identified in the brain^15,17–29^, including protofibrils which are targeted by lecanemab, a recently approved immunotherapy^30^.

Recently, in Liu *et al.* ^11^, we confirmed the key characteristics of Aβ*56 that were initially reported by Lesné *et al.*^16^, with the exception of its cellular compartmentalization. Liu *et al.*^11^ followed a protocol for compartmentalizing proteins that was briefly described in Lesné *et al.*^16^, with added details to compensate for missing information. Using this protocol, we demonstrated the presence of Aβ*56 in the extracellular space, confirming the original findings in Lesné *et al.*^16^. The replication and validation of the original findings in Lesné *et al.*^16^ are significant because Lesné *et al.*^16^ was recently retracted due to allegations of improper use of eraser and patch tools in WB’s depicted in Figure 2c and Supplementary Figure 4b. Comparing these two figures with the corresponding ones in the current paper ‒ specifically, Figure 2c in Lesné *et al.*^16^ with **Fig. 2b** and Supplementary Figure 4b in Lesné *et al.*^16^ with **Fig. 1e** ‒ it is possible that the improper alterations removed non-specific bands, but this could not be directly addressed because the corresponding source files were unavailable. It appears, however, that any potential inappropriate alterations in Lesné *et al.*^16^ did not involve manipulating relevant experimental signals. Notably, **Fig. 1e** in the current paper and Supplementary Figure 4b in Lesné *et al.*^16^ are identical within the limits of blot-to-blot variation. **Fig. 2b** in the current paper shows that Aβ*56 elutes in the expected fractions, indicating that it is not artificially generated through the aggregation of Aβ monomers due to exposure to SDS, thereby confirming the findings in Figure 2c of Lesné *et al.*^16^.

Retracting findings that have been independently replicated and confirmed may confound the scientific record by potentially misleading people into disbelieving accurate results. The confirmatory findings in Liu *et al.*^11^ and the current paper correct the scientific record and validate the biochemical and biological properties of Aβ*56 originally reported in Lesné *et al.*^16^.

## Acknowledgements

These studies were supported using funds from the N. Bud Grossman Center for Memory Research and Care and a Metropolitan Life Prize for Alzheimer’s Research awarded to K.H.A.

## Author contributions

P.L. conceived the study, designed and performed biochemistry experiments, analyzed data, and wrote the manuscript. I.P.L. helped design and perform biochemistry experiments, and analyzed data. K.H.A. conceived the study, analyzed data, and wrote the manuscript.

## Methods

### Mice

All experiments involving mice were performed in full accordance with the guidelines of the Association for Assessment and Accreditation of Laboratory Animal Care and approved (approval #1202A09927) by the Institutional Animal Care and Use Committee at the University of Minnesota, Twin Cities, Minnesota. Mice were conventionally housed in a vivarium and maintained on a 12-hr ON/12-hr OFF light cycle in which lights were on from 6 am to 6 pm. Mice were given *ad libitum* access to food and water.

Nine-month, female Tg2576 mice^31^ harboring the 695-amino-acid human amyloid precursor protein isoform with the Swedish (K670N, M671L) mutation (APP_695_-Swe) in C57B6/SJL background and age-matched, female, non-transgenic littermates were anesthetized using isoflurane and sacrificed by decapitation. For each animal, the brain was immediately harvested, and the olfactory bulb and cerebellum were then separated from the forebrain for each hemisphere. The hemi-forebrains of each mouse were collected, snap-frozen on dry ice, and stored at −80°C until further use. Basic information of all mice used is shown in Table 1.

**Table 1.**
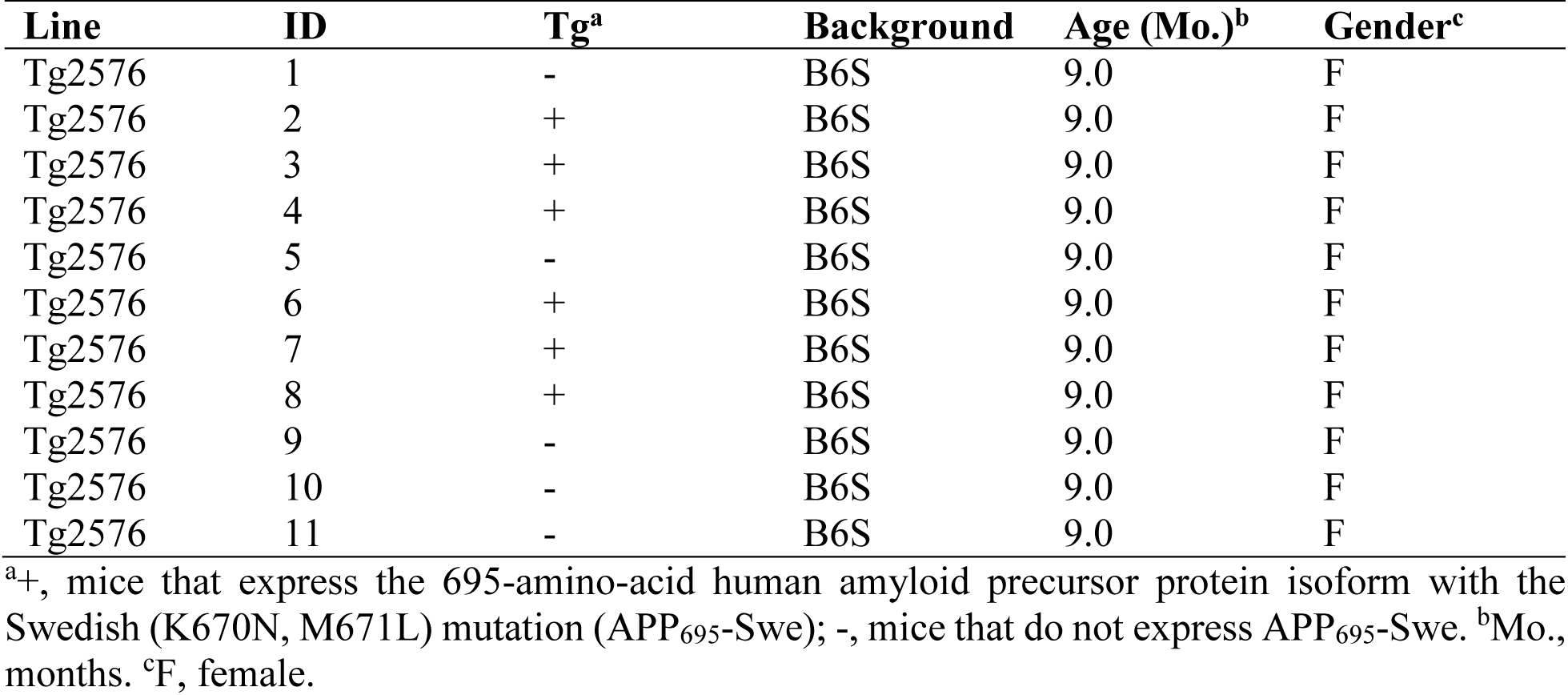
Mice used in this study.

| Line | ID | Tg<sup>a</sup> | Background | Age (Mo.)<sup>b</sup> | Gender<sup>c</sup> |
| --- | --- | --- | --- | --- | --- |
| Tg2576 | 1 | - | B6S | 9.0 | F |
| Tg2576 | 2 | + | B6S | 9.0 | F |
| Tg2576 | 3 | + | B6S | 9.0 | F |
| Tg2576 | 4 | + | B6S | 9.0 | F |
| Tg2576 | 5 | - | B6S | 9.0 | F |
| Tg2576 | 6 | + | B6S | 9.0 | F |
| Tg2576 | 7 | + | B6S | 9.0 | F |
| Tg2576 | 8 | + | B6S | 9.0 | F |
| Tg2576 | 9 | - | B6S | 9.0 | F |
| Tg2576 | 10 | - | B6S | 9.0 | F |
| Tg2576 | 11 | - | B6S | 9.0 | F |

<sup>a</sup>+, mice that express the 695-amino-acid human amyloid precursor protein isoform with the Swedish (K670N, M671L) mutation (APP<sub>695</sub>-Swe); -, mice that do not express APP<sub>695</sub>-Swe. <sup>b</sup>Mo., months. <sup>c</sup>F, female.

### Antibodies

Basic information of all antibodies used is shown in Table 2.

**Table 2.** Key resources used in this study.

| REAGENT or RESOURCE | SOURCE | IDENTIFIER |
| --- | --- | --- |
| **Antibodies** |  |  |
| Mouse monoclonal biotinylated anti-Aβ(3-8) | BioLegend | Cat# 803009; Clone# 6E10; Lot# B202110; RRID: AB_2564656 |
| Mouse monoclonal anti-Aβ(3-8) | BioLegend | Cat# 803003; Clone# 6E10; Lot# B353947; RRID: AB_2564652 |
| Mouse monoclonal biotinylated anti-Aβ(1-x) | IBL America | Cat# 10326; Clone# 82E1; Lot# 2K-115; RRID: AB_1540459 |
| Rabbit monoclonal anti-Aβ(x-40) | Cell Signaling Technology | Cat# 12990BF; Clone# D8Q7I; Lot# 2; RRID: AB_2798082 |
| Rabbit polyclonal biotinylated anti-amyloid oligomers | StressMarq Biosciences | Cat# SPC-506D; Clone# A11; Lot# MA333444; RRID: AB_10962958 |
| Mouse monoclonal anti-human sAPPα | IBL America | Cat# 11088; Clone# 2B3; Lot# 1I-227; RRID: AB_494690 |
| Rabbit polyclonal anti-C-terminus of human amyloid precursor protein | Thermo Fisher Scientific | Cat# 36-6900; Lot# QD218360; RRID: AB_2533275 |
| Rabbit polyclonal anti-C-terminus of human amyloid precursor protein | MilliporeSigma | Cat# A8717; Lot# 0000187789; RRID: AB_258409 |
| ImmunoPure goat anti-rabbit IgG (H + L) conjugated to horseradish peroxidase (HRP) | Thermo Fisher Scientific | Cat# 31463; Lot# LB12860121; RRID: AB_228333 |
| ImmunoPure goat anti-mouse IgG (Fc) conjugated to horseradish peroxidase (HRP) | Thermo Fisher Scientific | Cat# 31437; Lot# GF9459812; RRID: AB_228295 |
| **Biological samples** |  |  |
| Frozen forebrain tissue of Tg2576 mice (B6;SJL) | Dr. Karen Ashe laboratory | N/A |
| **Experimental models: Organisms/strains** |  |  |
| Mouse: Tg2576, B6;SJL-Tg(APP<sup>SWE</sup>)2576Kha/Tac: B6;SJL | Hsiao et al.<sup>31</sup> | RRID: IMSR_TAC:1349 |
| **Software and algorithms** |  |  |
| GraphPad Prism version 8.3.0 | GraphPad Software | RRID: SCR_002798 |
| Image Lab version 6.1 | Bio Rad | RRID: SCR_014210 |
| Microsoft PowerPoint 2016 | Microsoft | RRID: SCR_023631 |
| **Other** |  |  |
| Protein G Sepharose 4 Fast Flow resin | GE healthcare | Cat# GE17-0618-02 |
| Dynabeads Protein G | Thermo Fisher Scientific | Cat# 10004D |
| Neutravidin conjugated to horseradish peroxidase (HRP) | Thermo Fisher Scientific | Cat# A2664 |
| Superdex 75 Increase 10/300 GL column | Cytiva | Cat# 29-1487-21 |
| BioLogic DuoFlow Chromatography system | Bio Rad | RRID: SCR_019686 |
| ChemiDoc MP Imaging System | Bio Rad | RRID: SCR_019037 |

### Brain protein extraction

Protein extraction from forebrains were carried out following a protocol previously described.^16^ Specifically, each frozen hemi-forebrain free of olfactory bulb and cerebellum was transferred from −80°C directly to 0.5 mL of ice-cold extraction buffer 1 (50 mM tris(hydroxymethyl)aminomethane-hydrochloric acid (Tris-HCl), pH 7.6; 150 mM sodium chloride (NaCl); 2 mM ethylenediaminetetraacetic acid (EDTA); 0.01% (v/v) octyl phenoxypolyethoxylethanol (also known as nonidet P-40); 0.1% (w/v) sodium dodecyl sulfate (SDS); 0.1 mM phenylmethylsulfonyl fluoride (PMSF); 0.2 mM 1,10-phenanthroline monohydrate (phen); 1% (v/v) protease inhibitor cocktail (MilliporeSigma, Burlington, MA); 1% (v/v) phosphatase inhibitor cocktail A (SantaCruz Biotechnology, Dallas, TX); and 1% (v/v) phosphatase inhibitor cocktail 2 (MilliporeSigma)). The tissue was immediately homogenized at room temperature by first pulling the plunger of a 1-mL polypropylene syringe to draw it into the barrel through the tip. Next, a hypodermic needle (20-gauge x 1 inch) with aluminum hub was installed at the tip of the syringe, and the brain tissue was gently pushed to pass through the needle. The tissue was then pulled in and pushed out of the syringe for a total of 10 times. The time from the touch of the frozen tissue to the buffer to the completion of the 10-cycle homogenization was between 120 and 150 sec. The resulting brain homogenates from each mouse were immediately centrifuged at 4°C, 800 *g* for 5 min. Next, the supernatant was collected and immediately centrifuged at 4°C, 16,100 *g* for 1 min, and the pellet obtained from the 5-min centrifugation was placed briefly on ice. The supernatant of the 1-min centrifugation was collected and immediately centrifuged at 4°C, 16,100 *g* for 90 min. At the completion of the 90-min centrifugation, the supernatant was collected as the extracellular-enriched brain extracts (EC).

Immediately after the start of the 90-min centrifugation of the supernatant as described above, 0.5 mL of ice-cold extraction buffer 2 (50 mM Tris-HCl, pH 7.6; 150 mM NaCl; 0.1% (v/v) polyethylene glycol *p*-(1,1,3,3-tetramethylbutyl)-phenyl ether (Triton X-100); 0.1 mM PMSF; 0.2 mM phen; 1% (v/v) protease inhibitor cocktail; 1% (v/v) phosphatase inhibitor cocktail A; and 1% (v/v) phosphatase inhibitor cocktail 2) was added to the pellet that was stored on ice. The pellet and buffer 2 were mixed at room temperature by pipetting in and out of a 1,000-µL pipette tip for 10 cycles to achieve full resuspension. The resulting brain homogenates were immediately centrifuged at 4°C, 16,000 *g* for 90 min, and the supernatant was collected as the intracellular brain extracts (IC).

Immediately after obtaining the IC, 0.5 mL of ice-cold extraction buffer 3 (50 mM Tris-HCl, pH 7.4; 150 mM NaCl; 0.5% (v/v) Triton X-100; 1 mM ethylene glycol-bis(*β*-aminoethyl ether)-*N*,*N*,*N′*,*N′*-tetraacetic acid (EGTA); 3% (w/v) SDS; 1% (w/v) deoxycholate; 0.1 mM PMSF; 0.2 mM phen; 1% (v/v) protease inhibitor cocktail; 1% (v/v) phosphatase inhibitor cocktail A; and 1% (v/v) phosphatase inhibitor cocktail 2) was added to the pellet. The pellet and buffer 3 were mixed at room temperature by pipetting in and out of a 1,000-µL pipette tip for 20 cycles to achieve full resuspension. The resulting brain homogenates were rotated at 4°C, 15 rpm for 15 min, and then centrifuged at 4°C, 16,000 *g* for 90 min. The supernatant was collected as the membrane-bound brain extracts (MB).

Prior to biochemical analyses, each studied sample—the entire EC and entire MB prepared from each hemi-forebrain—was incubated with 40 µL of Protein G Sepharose 4 Fast Flow resin (GE healthcare, Piscataway, NJ) slurry (volume of settled beads:volume of storage buffer (50 mM Tris-HCl, pH 7.4; 150 mM NaCl) = 1:1) to deplete endogenous mouse immunoglobulin G (msIgG). Depletion of msIgG was carried out twice; each time, the mixture of the 40-µL slurry and the sample was rotated at 4°C, 15 rpm for 1 hr.

Protein concentrations were determined using the bicinchoninic acid (BCA) assay (Thermo Fisher Scientific, Waltham, MA). Brain extracts were stored at −80°C in multiple aliquots to preserve sample quality.

### Size-exclusion chromatography (SEC)

SEC was performed using a Superdex 75 Increase 10/300 GL column (Cytiva, Marlborough, MA) under the control of the BioLogic DuoFlow Chromatography system (Bio-Rad, Hercules, CA).

To calibrate the column, each entity in a set of biomolecule standards—four globular proteins at the molecular weights of 66, 29, 12.4, and 6.5 kDa, and Blue Dextran of >2,000 kDa (MilliporeSigma)—was diluted in elution buffer (50 mM 0.22 µm-filtered, autoclave-sterilized ammonium acetate, pH 8.5) and injected individually into the column at the dose of 200 µg. Biomolecule standards were eluted at room temperature with elution buffer at a flow rate of 0.3 mL/min and a fraction size of 500 µL. The eluted materials were collected into 1.5-mL microcentrifuge tubes for 48 fractions. The elution profile of each standard was determined using the BCA assay with a DTX880 Multimode detector (Beckman Coulter, Brea, CA).

To analyze biological samples, EC of female Tg2576 mice at 9 months of age before neuritic plaques emerge in the forebrain^32^ were pooled from hemi-forebrain extracts of 3 mice, each contributing 33% of the volume. The 500-µL injectates contained 2.8 mg of total proteins. Protein elution was carried out under the same experimental settings as described above except that 10 µL of elution buffer containing 5.1 mM PMSF, 10.2 mM phen, 5.1% (v/v) protease inhibitor cocktail, 5.1% (v/v) phosphatase inhibitor cocktail A, and 5.1% (v/v) phosphatase inhibitor cocktail 2 was added to each fraction-collecting microcentrifuge tube prior to the elution. In doing so, the post-elution concentrations of the five types of inhibitors were 0.1 mM, 0.2 mM, 0.1% (v/v), 0.1% (v/v), and 0.1% (v/v), respectively. Immediately after the elution was complete, the collected fractions were stored on ice. Each fraction was evenly split into two halves, snap-frozen on dry ice, and stored at −80°C. As a negative control, EC of hemi-forebrains of three age-matched, female, non-transgenic littermates of Tg2576 mice were prepared, pooled, and size-fractionated similarly as described above.

### Immunoprecipitation (IP)

To reveal the difference in expression pattern of fl-APP and its metabolites between the EC and MB, IP was performed based on previously published procedures^33^. Briefly, the EC and MB were incubated with the mouse monoclonal 6E10 antibody (BioLegend, San Diego, CA) that recognizes amino acids 3-8 of Aβ and the Protein G Sepharose 4 Fast Flow resin (GE healthcare) (**Tables 2** and **3**) by rotating at 4°C, 15 rpm for 14-16 hr. Subsequently, the immunocomplex-bound resin was first washed in 500 µL of wash buffer 1 (50 mM Tris-HCl, pH 7.4; 300 mM NaCl; 1 mM EDTA; 0.1% (v/v) Triton X-100; 0.1 mM PMSF; 0.2 mM phen; 0.1% (v/v) protease inhibitor cocktail; 0.1% (v/v) phosphatase inhibitor cocktail A; and 0.1% (v/v) phosphatase inhibitor cocktail 2) by rotating at 4°C, 15 rpm for 5 min, and then in wash buffer 2 (50 mM Tris-HCl, pH 7.4; 150 mM NaCl; 1 mM EDTA; 0.1% (v/v) Triton X-100; 0.1 mM PMSF; 0.2 mM phen; 0.1% (v/v) protease inhibitor cocktail; 0.1% (v/v) phosphatase inhibitor cocktail A; and 0.1% (v/v) phosphatase inhibitor cocktail 2) by rotating at 4°C, 15 rpm for 5 min. Detailed usage of brain protein extracts and reagents are listed in Table 3.

**Table 3.**
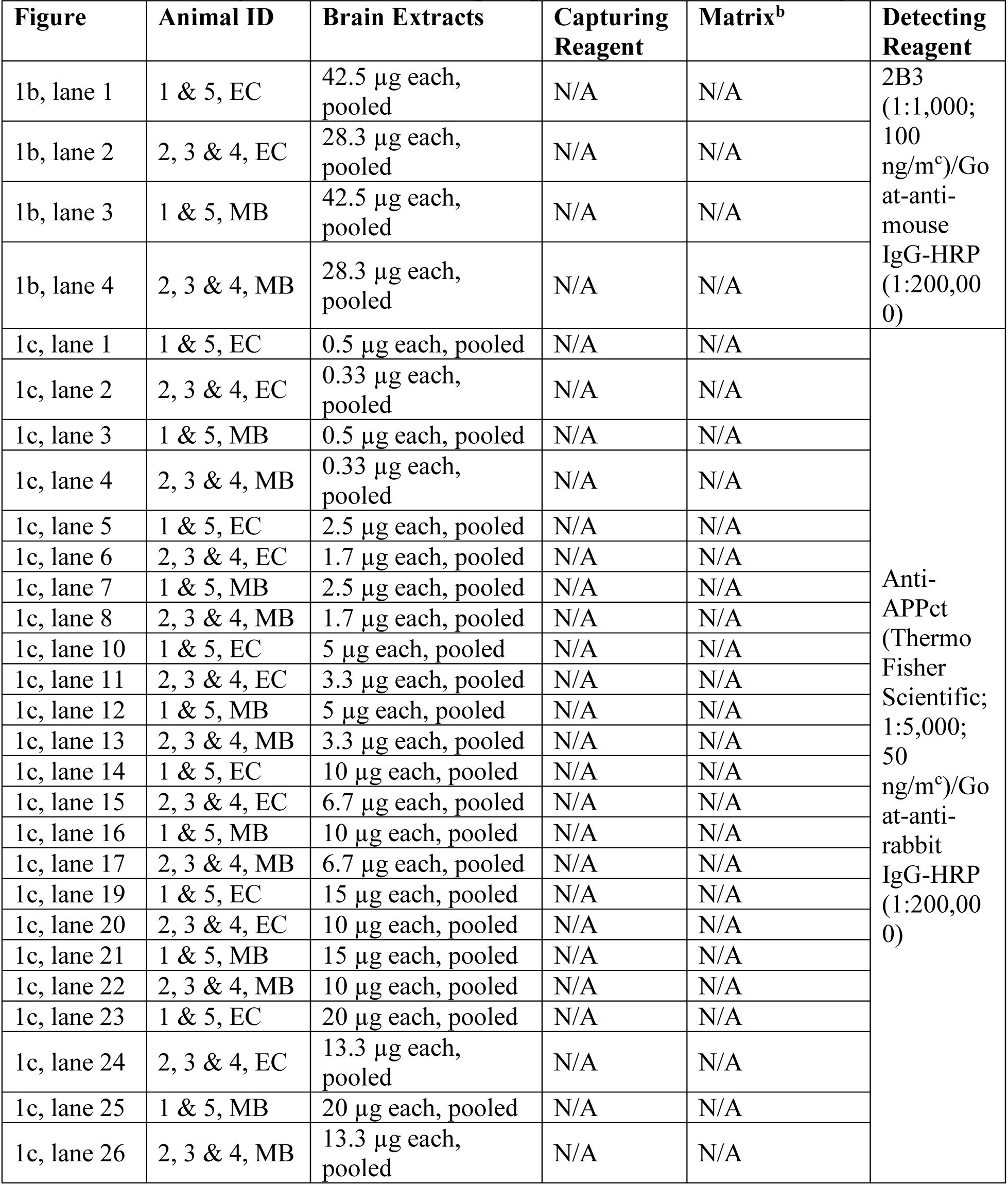
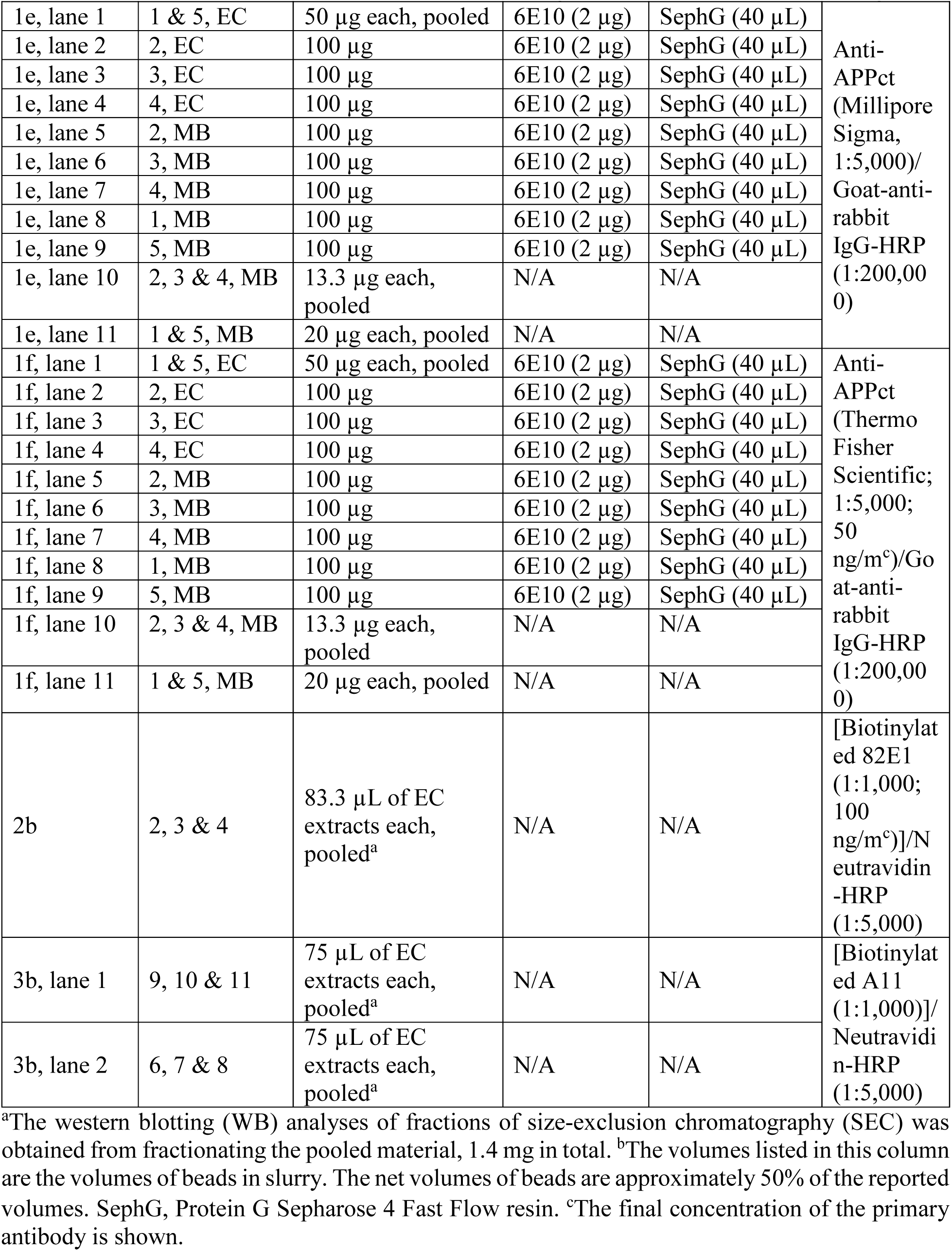
Mice and reagents used in immunoprecipitation and western blotting.

### Antibody immobilization on matrix

The antibody immobilization was carried out based on previous publications^11,34^. Briefly, 62 mg of the D8Q7I antibody that is directed against the C-terminus of Aβ(x-40) was incubated with 1 mL of Dynabeads Protein G matrix slurry (Thermo Fisher Scientific) at 4°C, 15 rpm for 14-16 hr. The antibody was crosslinked to the matrix by incubating with 70 mM dimethyl pimelimidate (Thermo Fisher Scientific; dissolved in 200 mM triethanolamine, pH 9.0 (MilliporeSigma)) solution at room temperature, 15 rpm for 15 min. Next, the chemical reaction was quenched by incubating the antibody-matrix complex with 10% (v/v) ethanolamine (MilliporeSigma) at room temperature, 15 rpm for 15 min. To eliminate non-covalently linked antibodies, beads were pre-treated with elution buffer 1 (100 mM glycine-HCl, pH 2.9; and 1 M urea) twice, each time at room temperature, 15 rpm for 5 min. The final product was stored at 4°C in TBS, pH 7.4, containing 0.05% (w/v) sodium azide and inhibitors (0.1 mM PMSF, 0.2 mM phen, 0.1% (v/v) protease inhibitor cocktail, 0.1% (v/v) phosphatase inhibitor cocktail A, and 0.1% (v/v) phosphatase inhibitor cocktail 2).

### Immunoaffinity purification

SEC fractions 22 and 23 of EC (**Tables 1-3**) of 9-month female Tg2576 or age- and gender-matched, non-transgenic mice were incubated with D8Q7I-bound Dynabeads matrix (**Tables 2** and **3**) at 4°C, 15 rpm for 20 hr. The immunocomplex-matrix was washed twice, each time with wash buffer 3 (TBS, pH 7.4; and 1% (w/v) *n*-Octyl *β*-*D*-thioglucopyranoside (OTG)) at 4°C, 15 rpm for 5 min. Immunoprecipitated proteins were eluted three times, each time with 100 µL of elution buffer 2 (Pierce IgG Elution Buffer, pH ∼3 (Thermo Fisher Scientific); 1% (w/v) OTG; and 2 M urea) at 50°C, 1,250 rpm for 5 min. Immediately after each elution, the eluted material was neutralized to pH 7-8 using 1 M Tris-base (pH ∼10.5). The eluted materials were snap-frozen on dry ice and stored at −80°C.

### Electro-elution

To fractionate the immunoaffinity-purified proteins by size, the eluted materials were subjected to SDS-PAGE using 1.0-mm-thick Novex 10–20% tricine gels under semi-denaturing conditions (samples in Laemmli sample buffer with no reducing agents or boiling treatment prior to electrophoresis). A portion (10%) of the fractionated proteins were then subjected to WB analyses probed with the biotinylated 82E1 antibody as described below.

Obtained WB images were used to identify the location of the ∼56-kDa entity and guide excising gel pieces that contained this molecule of interest from the other 90% of size-fractionated eluates. Next, the gel pieces were subjected to electro-elution using a Bio-Rad Model 422 Electro-eluter (Bio-Rad) as described in detail previously^11^. Briefly, electro-elution was performed at room temperature in elution buffer 4 (50 mM ammonium bicarbonate, pH 8.8; and 0.1% (w/v) SDS) under a constant current of 10 mA for 6 hr. The SDS detergent of the samples collected from electro-elution was removed at room temperature using the High Protein and Peptide Recovery Detergent Removal Resin (Thermo Fisher Scientific).

### Western blotting (WB)

WB was performed based on previously published procedures^7,33^. Detailed usage of brain protein extracts and detecting reagents are listed in Table 3.

Briefly, eluted materials from IP (**Figs. 1e** and **1f**), SEC-fractionated EC (**Fig. 2b**), EC and MB brain extracts (**Figs. 1b**, **1c**, **1e**, and **1f**), and immunoaffinity-purified/electro-eluted proteins (**Fig. 3b**) were electrophoretically separated by SDS-PAGE. To prepare eluted materials from IP, immunocomplexes (**Figs. 1e** and **1f**) were eluted by agitatedly heating the resin in 4x Laemmli sample buffer (Bio-Rad; in the presence of 1.42 M β-mercaptoethanol) at 95°C, 1,250 rpm for 10 min. To prepare loading samples for SDS-PAGE, SEC-fractionated EC (**Fig. 2b**), or EC and MB (**Figs. 1b**, **1c**, **1e,** and **1f**) were mixed with 4x Laemmli sample buffer and 1.42 M β-mercaptoethanol at 95°C, 1,250 rpm for 5 min. To prepare loading immunoaffinity-purified/electro-eluted proteins (**Fig. 3b**), samples were mixed with 4x Laemmli sample buffer at room temperature. Electrophoresis was performed at room temperature in running buffer 1 (Cathode buffer – 100 mM Tris-HCl, pH 8.25; 100 mM tricine; and 0.1% (w/v) SDS, and anode buffer – 200 mM Tris-HCl, pH 8.9) when 1.0-mm-thick, Novex 10-20% tricine gels (Thermo

Fisher Scientific) were used either under a voltage of 80 V for 180 min (**Figs. 1b**, **1e**, **1f**, and **2b**) or under a voltage of 125 V for 90 min (**Fig. 3b**). Alternatively, electrophoresis was performed in running buffer 2 (1x XT MES Running Buffer, pH 6.4; Bio-Rad) under a voltage of 80 V for 130 min when 1.0-mm-thick, 4-12% Criterion XT Bis-Tris protein gels (Bio-Rad) were used (**Fig. 1c**). Next, the size-fractionated proteins were transferred to 0.2-µm (pore size) nitrocellulose membranes (Bio-Rad) using transfer buffer (25 mM Tris-HCl, 192 mM glycine, and 10% (v/v) methanol, pH not adjusted) at 4°C under a current of 0.4 A for 4 hr.

Then, each protein-blotted membrane was soaked in 50 mL of phosphate-buffered saline (PBS, pH 7.4) of room temperature. The membrane was subject to microwave heating under full power for 25 sec and then 15 sec with a 4-min cooling time period at room temperature after each heating process.

After heat-induced antigen retrieval, membranes (**Figs. 1b**, **1c**, **1e**, **1f**, and **2b**) were incubated in blocking buffer 1 (10 mM Tris-HCl, pH 7.4; 200 mM NaCl; 0.1% (v/v) polyoxyethylene (20) sorbitan monolaurate (Tween-20); and 5% (w/v) bovine serum albumin (MilliporeSigma)) by rotating at room temperature, 70 rpm for 1 hr to block non-specific interactions with the detecting reagents.

To reveal the WB pattern of SEC-fractionated EC probed with secondary detecting reagents, membranes (bottom panels of **Fig. 2b**) were washed in wash buffer 1 (10 mM Tris-HCl, pH 7.4; 200 mM NaCl; and 0.1% (v/v) Tween-20) five times, each time at room temperature, 80 rpm for 5 min. Following the wash step, membranes were incubated with horseradish peroxidase (HRP)-conjugated Neutravidin (Neutravidin-HRP; Thermo Fisher Scientific; diluted 1:5,000 in wash buffer 1 of room temperature) by rotating at room temperature, 70 rpm for 30 min. After the incubation, membranes were washed in wash buffer 1 five times, each time at room temperature, 80 rpm for 5 min.

To reveal the WB pattern of SEC-fractionated EC probed with biotinylated 82E1 antibody (upper and middle panels of **Fig. 2b**), membranes were blocked using blocking buffer 1 as described above. The primary antibody (**Tables 2** and **3**) was then added to the blocking buffer 1, and the membranes were incubated by rotating at 4°C, 70 rpm for 14-16 hr. After primary antibody incubation, the membranes were washed with wash buffer 1 five times, each time at room temperature, 80 rpm for 5 min. Following the wash step, the membranes were incubated with Neutravidin-HRP and then washed as described above.

For WB analyses probed with non-biotinylated primary antibodies (**Figs. 1b**, **1c**, **1e**, and **1f**), membranes were first blocked using blocking buffer 1 as described above. Next, primary antibodies (**Tables 2** and **3**) were added to the blocking buffer 1, and the membranes were incubated by rotating at 4°C, 70 rpm for 14-16 hr. Following the wash step described above, the membranes were then incubated with either goat-anti-mouse IgG-HRP (diluted 1:200,000 in wash buffer 1 of room temperature) for 2B3 (**Fig. 1b**) or HRP-conjugated ImmunoPure goat-anti-rabbit IgG (Thermo Fisher Scientific; diluted 1:200,000 in wash buffer 1 of room temperature) for anti-APPct antibodies (**Figs. 1c**, **1e**, and **1f**) by rotating at room temperature, 70 rpm for 1 hr. After the incubation, membranes were washed in wash buffer 1 as described above.

For WB analyses probed with biotinylated A11 antibody (**Fig. 3b**), after heat-induced antigen retrieval, the membrane was first blocked in blocking buffer 2 (10 mM Tris-HCl, pH 7.4; 200 mM NaCl; 0.01% (v/v) Tween-20; and 5% (w/v) Carnation instant non-fat dry milk powder) by rotating at room temperature, 70 rpm for 1 hr, and then incubated with the primary antibody (**Tables 2** and **3**) in blocking buffer 2 by rotating at 4°C, 70 rpm for 14-16 hr. After that, the membranes were washed with wash buffer 2 (10 mM Tris-HCl, pH 7.4; 200 mM NaCl; and 0.01% (v/v) Tween-20) five times, each time at room temperature, 80 rpm for 5 min. Following the wash step, the membrane was incubated with Neutravidin-HRP (diluted 1:5,000 in wash buffer 2 of room temperature) by rotating at room temperature, 70 rpm for 30 min. After the incubation, the membrane was washed in wash buffer 2 as described above.

For signal detection, membranes were incubated with either the SuperSignal West Pico PLUS Chemiluminescent Substrate (Thermo Fisher Scientific) by rotating at room temperature, 200 rpm for 4 min (**Figs. 1b** and **1f**) or 30 sec (**Fig. 1e**) or the SuperSignal West Femto Maximum Sensitivity Substrate (Thermo Fisher Scientific) at room temperature, 200 rpm for 5 min (**Figs. 1c**, **2b**, and **3b**). Chemiluminescence signals were developed using the ChemiDoc MP Imaging System (Bio-Rad). Densitometry-based quantification of protein band intensities (**Fig. 2c**) was performed using Image Lab version 6.1 (Bio-Rad).

The experimental settings for each (IP)/WB reaction are shown in Table 3.

